# Genomic Annotation Infrastructure (GAIn): Pipelines and Resource Repositories for Annotating Variants, Positions, and Regions

**DOI:** 10.64898/2026.07.08.737273

**Authors:** Murat Cokol, Lubomir Chorbadjiev, Yoon-ha Lee, Minal Jamsandekar, Ilina Gergova, Ivo Todorov, Ivan Iossifov

## Abstract

Interpretation of genomic variants, positions, and regions depends on reliable annotation—adding evidence such as predicted effect, conservation, population frequency, and gene-level context—yet the underlying resources are numerous, versioned, and assembly-specific. We present the Genomic Annotation Infrastructure (GAIn), a platform that generates transparent, reproducible annotations via declarative pipelines that define annotation tasks as ordered lists of components, called annotators, that produce annotation attributes using genomic resources from Genomic Resource Repositories (GRRs). We provide two public GRRs: a main repository containing more than 250 heterogeneous genomic resources, and a separate GRR-ENCODE repository containing resources derived from thousands of ENCODE (Encyclopedia of DNA Elements) project experiments. Users can use the annotation pipelines we made available, author custom annotation pipelines, and execute annotation tasks with these pipelines via GAIn’s web and command-line interfaces. The web interface can be used without any setup, but it relies on shared computational infrastructure and imposes limits on the size of annotation tasks. The command-line interface requires setup but supports arbitrarily large annotation tasks through simple-to-use parallelization and offers a broader set of features. For example, command-line GAIn can be extended by using custom GRRs or creating custom annotators via its plugin architecture. In addition, GAIn’s re-annotation feature, which updates annotations as they evolve, substantially simplifies maintaining annotations in a large genomics analysis project. GAIn’s resource management, explicit versioning, and pipeline abstraction provide an auditable, maintainable, and efficient foundation for modern genomic annotation across reference assemblies and use cases.

## Introduction

High-throughput sequencing has made it routine and increasingly inexpensive to sequence individual genomes, but every new genome brings a daunting number of variants to interpret^1–3^. A typical whole-exome sequence yields tens of thousands of single-nucleotide variants and indels per person, and a whole genome reveals millions of differences relative to the reference^4–6^. Only a small fraction of these variants is likely to contribute to disease risk, treatment response, or other phenotypes. The rest reflect benign population variation or neutral drift^7,8^. The falling cost of genomic assays has also enabled diverse experiments that generate large sets of genomic positions, such as GWAS hits or QTLs, and genomic regions, such as ChIP-seq or ATAC-seq peaks, that must likewise be interpreted and prioritized^9–11^.

Annotation is what turns genomic data into interpretable hypotheses, enabling clinicians and researchers to prioritize a handful of credible candidates from an otherwise overwhelming background of genomic variation^12,13^. Numerous groups have developed prediction scores and curated datasets that quantify different aspects of genomic variants, positions, or regions. Reference genomes and gene models provide essential context for interpretation, and other resource types contribute orthogonal information, including evolutionary conservation (e.g., phyloP^14^, phastCons^15^), functional constraint (e.g., FitCons^16^, LINSIGHT^17^), population frequency (e.g., gnomAD^18^), and curated clinical evidence (e.g., ClinVar^19^).

To make this growing ecosystem of annotations more accessible, several general-purpose genomic annotation frameworks were developed, though most are designed primarily for variant interpretation. ANNOVAR was among the first widely used tools to combine gene-based consequences, regional overlaps, population frequencies, and custom annotation sources in a single command-line workflow^20,21^. The Ensembl Variant Effect Predictor (VEP) provided tight integration with Ensembl gene models and regulatory annotations while also supporting many external scores^22^. More recently, OpenCRAVAT introduced a modular, plugin-based system with both graphical and programmatic interfaces^23,24^. These frameworks demonstrate that unifying heterogeneous resources behind a single interface greatly simplifies genomic annotation. However, the diversity of formats, coordinate systems, assembly, and gene model dependencies still calls for approaches that systematically organize resources, record provenance and versions, and let users specify complex annotation workflows in a transparent, reusable way.

Here we present the Genomic Annotation Infrastructure (GAIn), a framework designed to make genomic annotation transparent, modular, and reproducible in the face of this growing resource landscape. GAIn combines Genomic Resource Repositories (GRRs), which organize heterogeneous genomic resources with recorded provenance, with declarative annotation pipelines that specify how those resources are applied to each genomic variant, position, or region. We compiled and made publicly available two large GRRs: the main GRR, comprising more than 250 heterogeneous resources, and GRR-ENCODE, comprising resources from the nearly 8,000 ENCODE project experiments. Users can run pipelines at scale, annotating hundreds of millions of variants, from the command line or interactively via a web interface, using either our publicly available GRRs or their own private repositories that include additional or proprietary resources. Pipelines are expressed in a concise YAML format, making it straightforward to define, share, and version-control complex combinations of annotations. A plugin architecture and Python interface allow new annotators to be added without changing the core system, and re-annotation lets users update specific outputs when underlying resources change. Together, these features make GAIn a flexible platform for building and maintaining modern, scalable genomic annotation workflows.

## Results

### Overview

The main task of the Genomic Annotation Infrastructure (GAIn) is to assign annotation attributes to genomic objects, which we call annotatables. GAIn supports three types of annotatables: variants, positions, or regions. Figure 1 summarizes the main components of GAIn. Users specify the desired attributes in annotation pipelines, which are ordered lists of annotators. Annotators use diverse genomic resources, such as reference genomes, gene models, and position scores, to generate annotation attribute values, such as predicted functional effects and conservation scores. Annotators can also use outputs from earlier annotators within the same pipeline. Genomic resources are organized within a Genomic Resource Repository (GRR). We maintain two publicly available GRRs: our main GRR (https://grr.iossifovlab.com/) currently containing 272 resources, and GRR-ENCODE (https://grr-encode.iossifovlab.com/) containing resources derived from 7,922 ENCODE project experiments^10,25^. Users may also create their own custom GRRs to include additional resources.

**Figure 1.**
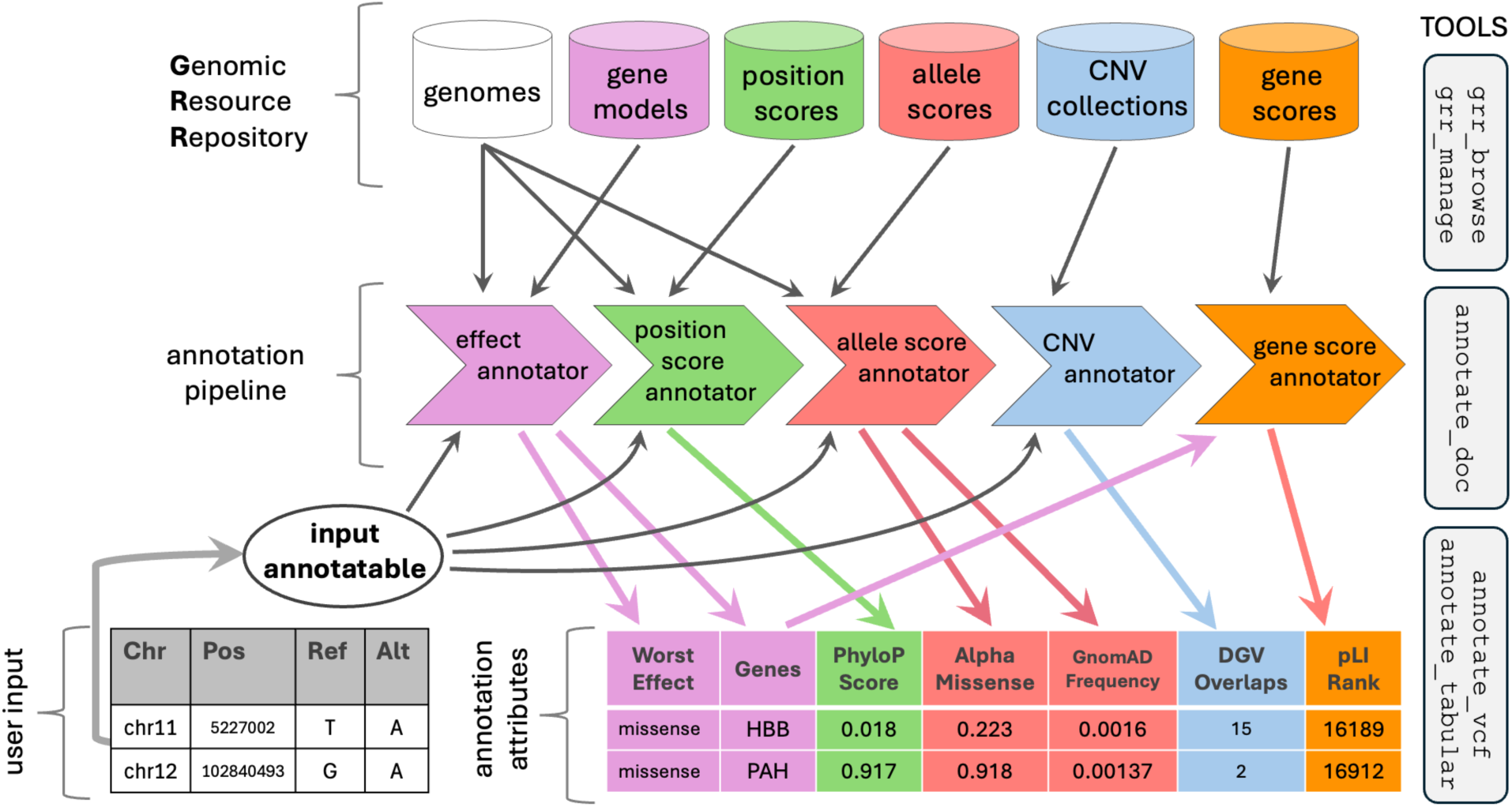
Overview of GAIn: resources, annotators, tools, and example workflow. (Top) Genomic resources, such as reference genomes, gene models, position scores, allele scores, CNV collections, and gene scores, are organized in a Genomic Resource Repository (GRR). **(Middle)** An annotation pipeline chains annotators: software components that produce annotation attributes. The pipeline here uses an effect annotator, describing the effect of the annotatable on genes, a position score annotator, assigning attributes that depend on chromosomal location, two allele score annotators, assigning attributes defined for specific alleles (e.g., pathogenicity or frequency), a CNV annotator, characterizing CNVs overlapping with the annotatable, and a gene-score annotator assigning attributes based on the genes implicated by the annotatable. Arrows indicate the resources each annotator uses; colors match resource and annotation attributes. Importantly, the gene score annotator takes the gene list produced by the effect annotator as input, demonstrating that annotators can use attributes produced earlier in the pipeline. **(Bottom)** The inputs for the annotation process can be table-like files (e.g., comma or tab-separated text files), as in this example, or VCF files. The two variants in this example correspond to mutations known to cause sickle cell disease and phenylketonuria. GAIn converts each row from the input file into an “input annotatable,” which is then passed to annotators. The resulting annotation attributes are added as columns to table-like input files (as shown in the figure) or as INFO field attributes to VCF input files. **(Right)** The main GAIn tools for exploring and managing a GRR (grr_browse, grr_manage), generating pipeline documentation (annotate_doc), and performing annotation (annotate_vcf, annotate_tabular).

GAIn provides a web interface (https://gain.iossifovlab.com/) that allows users to create custom annotation pipelines using the resources in our public GRRs. These custom pipelines, as well as pipelines included in our main GRR, can then be applied to large numbers of annotatables from tabular or VCF files. The full functionality of GAIn is accessible through its command-line tools, which enable users to efficiently annotate large numbers of variants, positions, or regions, build their own GRRs, browse available resources within a GRR, generate documentation for annotation pipelines, use the re-annotation features, and extend GAIn by authoring custom plug-in annotators. In the following sections, we describe the types of genomic resources used in GRR, demonstrate the use of annotators, and illustrate how to construct annotation pipelines. A complete description of GAIn is available in its documentation (https://iossifovlab.com/gaindocs).

### GAIn Components

#### Genomic Resource Types

GAIn utilizes multiple resource types (Figure 2). A complete list of supported resource types, along with detailed usage instructions, is available in the GAIn documentation. GAIn’s plugin architecture (see below) can be used to extend the platform with new resource types. Below, we describe GAIn’s core resource types.

**Figure 2.**
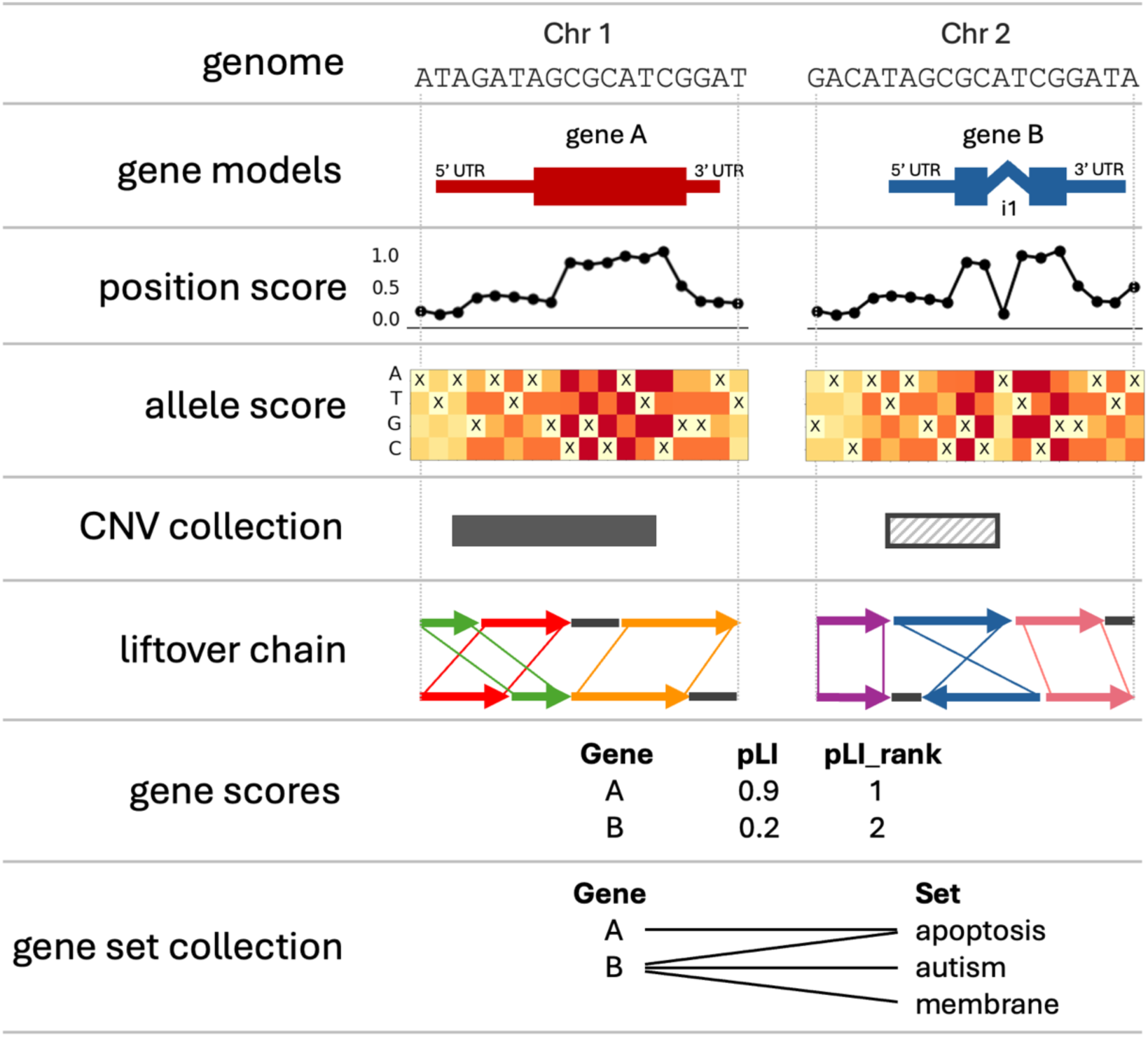
Genomic resource types supported by GAIn. Genomic resource types supported by GAIn are illustrated using a mock genome with two chromosomes. Resources of type “genome” contain reference genome sequence assemblies as indexed FASTA files. “Gene models” resources provide the location and structure of genes. In the small example, chromosome Chr1 contains gene A, and Chr2 contains gene B, which has an intron. “Position score” resources provide one or more scores for specific genomic locations (e.g., conservation). In this schematic, the coding regions score highest, UTRs score lower, and intronic and intergenic regions score lowest. “Allele score” resources provide information about specific non-reference variants (e.g., pathogenicity predictions or allele frequency). Here, the heatmaps indicate single-nucleotide variations and their hypothetical scores. Cells marked with ‘X’ correspond to the reference (no change) and are not scored, while substitutions are color-scaled to reflect low-to-high scores. “CNV collection” resources include observed duplications (e.g., on Chr1) or deletions (e.g., on Chr2). “Liftover chain” resources allow mapping genomic coordinates in one assembly to another, as shown here for our two-chromosome genome example. “Gene score” resources provide information about the genes implicated by the annotatable. Here, gene A has a high pLI score, indicating it is intolerant to mutations. “Gene set collection” resources organize genes into curated functional, pathway, or disease categories, providing functional context for each gene. Here, gene A is in the apoptosis set, and gene B is in the apoptosis, autism, and membrane sets.

**Genome** resources specify the reference genome assembly used to interpret all coordinates and alleles. In GAIn, this is an indexed FASTA and an optional list of chromosome/contig aliases (e.g., chr1/1). Widely used human reference builds include GRCh37/hg19 and GRCh38/hg38, which remain standards in clinical and research pipelines^26,27^. Recent long-read efforts have produced gap-free assemblies, such as T2T-CHM13/hs1, resolving centromeres and other previously missing regions^28^.

A “**gene models**” resource defines a set of transcripts for a specific reference assembly. Each transcript is characterized by its exon–intron structure, UTRs, coding sequences, splice junctions, strand, biotype, and stable identifiers, such as transcript ID and gene name. Gene models provide the structural context for interpreting variant effects. In GAIn, these resources may be in GTF or tabular formats, such as those used in the UCSC genome browser^29^. Historically, human annotation has centered on RefSeq^30^ and GENCODE^31^ gene models, which include many known transcripts per gene. By contrast, the MANE project selected a single, high-quality, representative transcript for each gene to provide a standard for clinical reporting and research^32^.

**Position score** resources assign per-nucleotide metrics independent of a particular allele, providing locus-level context such as evolutionary conservation or functional constraint. GAIn supports BigWig, indexed BedGraph, and tabix-indexed tabular files for this resource type. Examples include phyloP^14^, which measures evolutionary rate at the nucleotide level, and an ENCODE ChIP-seq experiment^25^ targeting a specific transcription factor in a specific tissue, showing the transcription factor’s binding level at every genomic position.

**Allele score** resources store information at the level of the variant allele, such as population-specific allele frequencies (e.g., gnomAD^18^) or pathogenicity predictions (e.g., CADD^33^, AlphaMissense^34^, MPC^35^). GAIn supports VCF and tabix-indexed tabular files for this resource type. Accurate use of allele resources requires canonical allele representation so that lookups resolve to the intended record^36,37^. Three specific allele-score resources included in our GRR are particularly noteworthy. dbNSFP^38^ is a consolidated variant database that aggregates many predictor outputs and related annotations into one lookup table that includes all possible non-synonymous single-nucleotide variants. ClinVar^19^ contributes curated clinical assertions (e.g., clinical significance labels and disease terms) for specific alleles, with provenance and review status. dbSNP provides stable identifiers (rsIDs) for known variants^39^.

**CNV collection** resources summarize previously observed copy-number variants (CNVs), including deletions or duplications of genomic segments, ranging from ∼50 bp to several megabases. In GAIn, CNV resources are represented as tabular files. Here we highlight four of the CNV collections included in the main GRR. The Database of Genomic Variants (DGV) aggregates structural variation from control populations^40^. dbVar is a public archive of submitted structural variants from research and clinical studies^41^. The gnomAD Exome CNV provides WES-based CNV calls in large cohorts^42^, while gnomAD Genome SV catalogs WGS-based structural variants, including CNVs, inversions, and complex SVs^43^. Both gnomAD collections include population frequency for each variant.

**Liftover chain** resources define directional mappings from a source to a target assembly needed to translate coordinates and associated alleles between references (e.g., from T2T-CHM13/hs1 to the GRCh38/hg38 reference genome)^44^. Because position, allele, and CNV resources are assembly-specific (e.g., phyloP7^14^ on hg38), liftover chains enable the use of these resources when the annotatables are defined on another assembly. GAIn supports liftover mappings with standard UCSC chain files.

**Gene score** resources are per-gene metrics that summarize gene properties such as tolerance to damaging mutations, haploinsufficiency, and coding length. After an annotatable is mapped to one or more genes by a gene model, these scores provide gene-level context for interpretation. In GAIn, gene scores are represented as tabular files keyed by gene names. For example, the LOEUF score reflects a gene’s intolerance to loss-of-function variants in the human population^18^, and the SFARI Gene score reflects the strength of the gene’s association with autism^45^.

**Gene set collection** resources group genes by shared functions, pathways, associated phenotypes, or other properties. Gene sets are built from expert-curated pathway/ontology knowledge or from experimental data, derived from expression, perturbation, or other experiments. For example, Gene Ontology resource^46^ uses expert curation to label genes (thus define gene sets) with terms from its three ontologies (Cellular Component, Molecular Function, and Biological Process), while the sets “targets of the RNA-binding protein FMRP”^47^ or “targets of the CHD8 chromatin modifier”^48^ are identified using experiments. In GAIn, gene sets are represented in GMT (Gene Matrix Transposed)^49^ or custom text formats.

#### Genomic Resource Repository (GRR)

Over the past few decades, researchers have produced a vast number of genomic resources, but these vary in assembly, format, version, and dependencies, complicating genomic annotation. GAIn addresses this challenge through Genomic Resource Repositories (GRRs): curated collections containing resources retrieved from their original sources, validated for integrity, harmonized into a common schema, and then version-pinned and indexed for fast, reproducible access. Each resource in a GRR is accompanied by an information page detailing its contents, sources, and annotation attributes. Users can either build private GRRs (described below) or point their annotation pipelines to our curated and actively maintained public GRRs, which aggregate widely used resources with recorded provenance (grr.iossifovlab.com and grr-encode.iossifovlab.com).

Our main public GRR currently comprises 272 resources organized by genome assembly: hg19, hg38, and hs1 (Figure 3a). The hg19 collection includes 2 genomes, 5 gene models resources, 135 position-scores (e.g., phastCons^15^, phyloP^14^, and multiple tissue-specific FitCons2^16^ tracks), and 4 allele-level resources (e.g., gnomAD^18^, CADD^33^, and MPC^35^). The hg38 collection is more extensive, with 4 genome resources, 51 gene models resources (e.g., multiple versions of MANE^32^ and GENCODE^31^), 8 position-score resources (e.g., phastCons^15^, phyloP^14^), 26 allele-score resources (e.g., gnomAD^18^, AlphaMissense^34^, ClinVar^19^, dbNSFP^38^, dbSNP^39^), and 6 CNV collection resources (e.g., DGV^40^, and dbVar^41^). For hs1, public position, allele, and CNV resources remain limited, and the GRR currently includes only 1 genome and 1 gene models resource. To bridge this gap, the public GRR also hosts liftover chain resources for mapping coordinates among hg19, hg38, and hs1 assemblies, enabling resources developed for one assembly to be applied to another. Importantly, many resources provide multiple scores; for example, the gnomAD resources include four scores, including alternate allele count, alternate allele frequency, and dbNSFP includes more than 400. Beyond assembly-specific resources, the main GRR provides 10 gene score resources (e.g., pLI^50^, RVIS^51^, LOEUF^18^, and autism gene scores; Figure 3b), and 8 gene set resources (e.g., GO^46^, MSigDB^52^, protein domains^53^) (Figure 3c).

**Figure 3.**
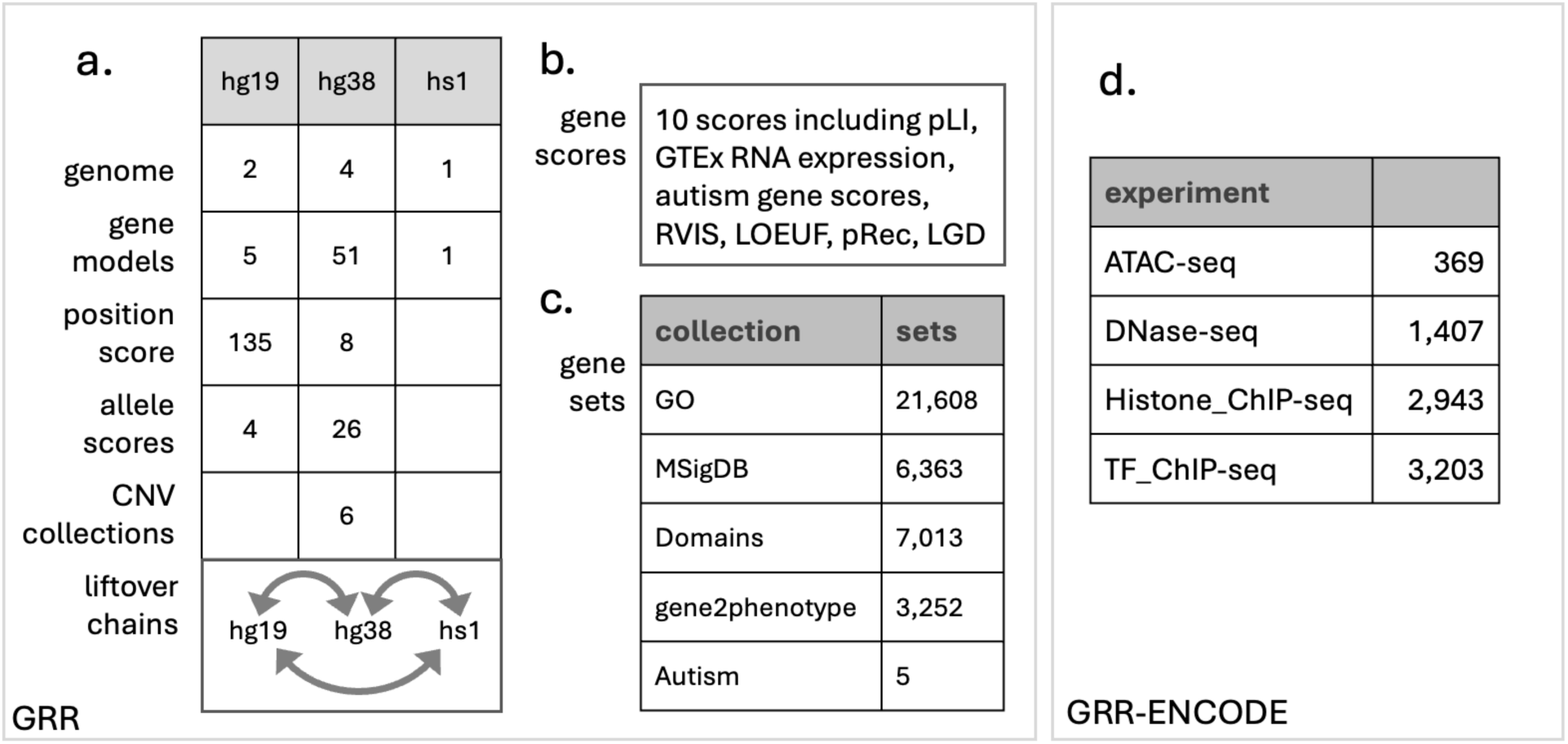
**Contents of our two public Genomic Resource Repositories (GRR)**. **(a)** Currently, our main GRR (https://grr.iossifovlab.com) includes resources for three assemblies: hg19, hg38, and hs1, along with several gene models resources for each assembly. Many position scores and allele scores are available for hg19 and hg38, and six CNV collections specific to hg38 are included. Our public GRR contains liftover chains, enabling mapping among the three assemblies. Our public GRR also contains **(b)** 10 gene scores covering various gene properties and **(c)** 8 gene set collections. **(d)** GRR-ENCODE (grr-encode.iossifovlab.com) contains 7,922 position scores in hg38 coordinates of four types: ATAC-seq (369), DNase-seq (1,407), Histone ChIP-seq (2,943), and TF ChIP-seq (3,203). Each ENCODE position score represents the result of the peak-calling pipelines applied to the corresponding ENCODE assay data.

To complement the main GRR, we created GRR-ENCODE, a separate public GRR dedicated to ENCODE-derived resources^10^ for use in GAIn annotation workflows (Figure 3d). All resources in GRR-ENCODE are position-score resources. Because ENCODE contributes many thousands of experiment-specific datasets, we maintain them in a dedicated repository rather than merging them into the main GRR. This separation preserves the browsability and heterogeneous character of the main repository while providing organized access to the full ENCODE collection. The current GRR-ENCODE release includes 369 ATAC-seq, 1,407 DNase-seq, 2,943 Histone ChIP-seq, and 3,203 TF ChIP-seq resources, all packaged in the same curated, versioned, and documented framework used throughout GAIn. This consistency allows users to inspect resource summaries and incorporate ENCODE-derived annotations reproducibly into their pipelines.

#### Annotatables and Annotation Pipelines

GAIn assigns attributes to objects called **annotatables**. GAIn supports three annotatable types: 1) a genetic variant, defined by its chromosome, position, reference, and alternative alleles; 2) a genomic position, defined by its chromosome and position, and 3) a genomic region, defined by its chromosome, start, and end coordinates. GAIn supports two input file formats. Tabular files (e.g., comma or tab-separated text file) contain one annotatable per line; the annotatable type is controlled by settings in the web interface or by command-line parameters. For tabular inputs, GAIn outputs a tabular file that preserves the input file columns and appends additional columns for the requested annotation attributes. GAIn can also annotate variant annotatables represented in VCF files (including multiple alternative allele lines), generating output VCFs with the requested annotation attributes stored as INFO fields.

An annotation pipeline is an ordered list of annotators that generate attributes for each annotatable, typically using one or more genomic resources from a Genomic Resource Repository (GRR). Annotators can also use attributes generated by earlier annotators upstream in the annotation pipeline. An annotation pipeline operates on a single annotatable at a time, which we call the “input annotatable”. By default, the “input annotatable” is available to every annotator in the pipeline. However, GAIn includes annotators that can produce derived annotatables (e.g., “liftover_annotator”, see below). Downstream annotators can then use either the “input annotatable” or a previously created annotatable (Figure 4).

**Figure 4.**
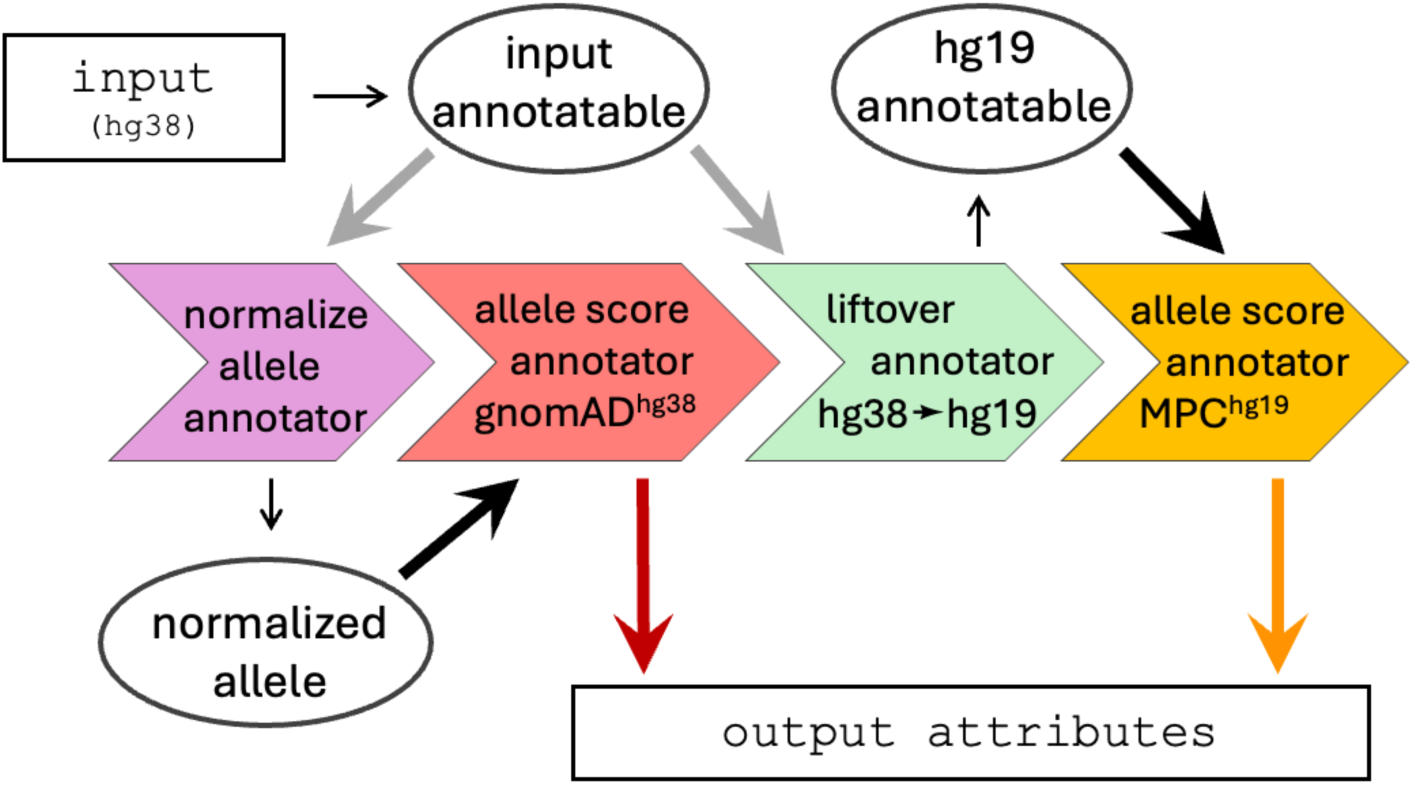
**Annotatables used during the execution of a GAIn pipeline**. The diagram shows three annotatable objects: the “input annotatable” (hg38), a derived “normalized allele” from “normalize_allele_annotator”, and a derived “hg19 annotatable” from “liftover_annotator”. Gray thick arrows indicate that the hg38 “input annotatable” is provided by default to the “normalize_allele_annotator” and the “liftover_annotator”. The “normalize_allele_annotator” canonicalizes the hg38 allele (e.g., left-normalizes indels), enabling direct lookup of allele frequencies in resources (e.g., gnomAD) that use canonical alleles. The “liftover_annotator” converts the annotatable to hg19 coordinates, producing an “hg19 annotatable” that is explicitly (dark thick arrows) passed to an allele score annotator, which uses the hg19-defined MPC resource (MPC). Downward arrows mark the emitted attributes: gnomAD outputs allele frequency (red), and MPC outputs a missense impact score (orange).

Annotation pipelines are YAML^54^ files with a specific structure that includes an optional preamble and a list of annotators (see Figure 5). For each annotator, the user defines an annotator type, provides the IDs of the required resources, selects the annotatable (“input_annotatable” is used by default), sets additional configuration parameters as needed (e.g., “promoter_len” for “effect_annotator”), and selects attributes for the output. If no attributes are explicitly specified, the annotator emits its default attribute set, which may depend on the selected resources. The user can further configure explicitly selected attributes, such as renaming them or selecting value aggregators. For example, annotator 4 in Figure 5, which is of type “allele_score_annotator”, will use the resource with id “hg38/variant_frequencies/gnomAD_4.1.0/exomes/ALL” and the non-default annotatable called “normalized_allele”, and will output the AF score from the gnomAD resource as an attribute named “gnomAD_4.1_af”.

**Figure 5.**
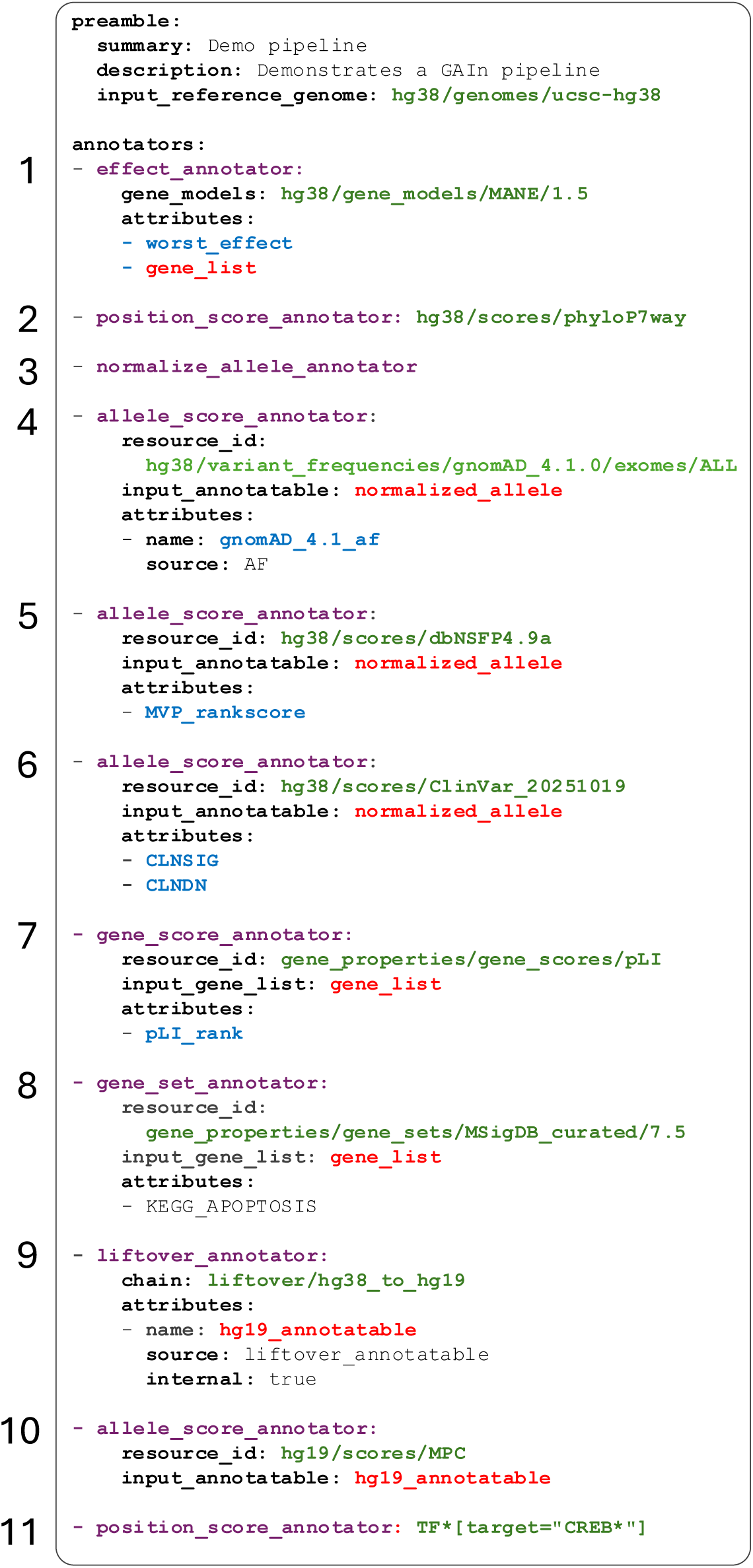
Example GAIn pipeline generating multiple annotations. The optional preamble provides a summary and description of the pipeline and sets the reference genome (hg38) expected for the pipeline inputs. (1) “effect_annotator” uses the selected gene model (MANE 1.5) to identify implicated genes (gene_list) and the worst predicted consequence (worst_effect) across all transcripts that overlap the variant. (2) “position_score_annotator” queries the selected resource (phyloP7way) at the variant’s coordinate and outputs the phyloP7way score. (3) “normalize_allele_annotator” creates a “normalized_allele” annotatable for downstream allele score annotators. (4) “allele_score_annotator” uses the gnomAD v4.1.0 resource on the “normalized_allele” annotatable to look up the alternative allele frequency (AF) and outputs it as gnomAD_4.1_af. (5–6) Two “allele_score_annotators” run on the “normalized_allele” annotatable: one queries dbNSFP v4.9a to return MVP_rankscore (rank score for predicted missense deleteriousness), one of its more than 400 scores; the other uses ClinVar to return CLNSIG (ClinVar clinical significance label) and CLNDN (ClinVar disease name). (7) “gene_score_annotator” uses the pLI gene resource with the gene_list produced by the effect annotator in (1) and outputs pLI_rank (loss-of-function intolerance rank). (8) “gene_set_annotator” uses MSigDB_curated 7.5 gene set collection resource with the gene_list and outputs a Boolean attribute indicating membership in ‘KEGG_APOPTOSIS’. (9) “liftover_annotator” (hg38→hg19) applies the liftover/hg38_to_hg19 chain to create an internal hg19_annotatable, for use by hg19-specific resources, that is labeled as ‘internal’, meaning it is not emitted in the output but is only used by downstream annotators. (10) “allele_score_annotator” uses the hg19/MPC resource with the hg19_annotatable and adds the corresponding MPC score. (11) A shorthand annotator definition expands into multiple “position_score_annotators”, one for each ENCODE resource matching the pattern “TF*[target=CREB*]”, or transcription-factor ChIP-seq experiments that target proteins whose names start with “CREB.”

GAIn supports many annotators out of the box and allows creating new annotators through its plugin architecture (see below). GAIn’s documentation includes an up-to-date list of all supported annotators, along with detailed descriptions of their configuration parameters and the attributes they produce. Below, we will briefly describe the most used annotators.

“**effect_annotator**” identifies the genes affected by the annotatable and, for variant annotatables, predicts their effects on the encoding of the affected genes using a gene models resource. GAIn’s “effect_annotator” is optimized for annotation of genetic variants and can produce more than 20 attributes (e.g., transcript-level effect details, lists of noncoding genes).

**“simple_effect_annotator”** also identifies the genes affected by the annotatable but uses a simple-effect classification we developed based on the types of transcript elements that overlap the genomic region associated with the processed annotatable (see Figure 2 in ^55^). The “simple_effect_annotator” outputs attributes like those of the “effect_annotator,” but is optimized for use with genomic position or genomic region input annotatables.

**“position_score_annotator”** uses the scores from the selected position score resource to assign attributes to the annotatable. When the annotatable covers multiple genomic positions (e.g., short deletions or genomic region annotatables), the annotator uses an aggregator (e.g., “max”, “mean”, or “list”) to combine the scores for each of the covered positions.

**“allele_score_annotator”** adds attributes to annotatables using scores from a given allele score resource. The annotator matches a variant annotatable to the alleles in the resource and copies the scores as attributes, or, for region annotatables, identifies alleles in the region and aggregates their score values.

**“gene_score_annotator”** adds gene-level metrics for genes implicated by the annotatable using gene score resources. It takes an “input_gene_list” specifying a gene list attribute, which is usually generated by effect or simple-effect annotators earlier in the pipeline. When multiple genes are affected by the annotatable, the “gene_score_annotator” aggregates the scores across all genes.

**“gene_set_annotator”** adds gene-set membership information using gene-set resources. Like the “gene_score_annotator”, it operates on a list of genes generated from previous annotators. It emits true/false attributes based on whether the input genes overlap with specified gene sets from the gene-set resource. Alternatively, the annotator can generate a list of all gene sets from the resource that the input genes belong to.

**“cnv_collection_annotator”** queries CNV collection resources to report overlap between the annotatable and previously observed deletions/duplications, optionally restricted by user-defined filters (e.g., CNV type, size, or frequency).

**“normalize_allele_annotator”** converts a variant to the canonical allele representation and produces a “normalized_allele” annotatable, which downstream annotators (e.g., “allele_score_annotator”) can use.

**“liftover_annotator”** maps genomic coordinates between assemblies by using liftover chain resources, and produces “liftover_annotatable”, which downstream annotators use.

Figure 5 shows an example GAIn pipeline with multiple annotators and a preamble. The preamble provides a summary and description of the pipeline and sets the default genome to “hg38/genomes/ucsc-hg38”, indicating that the pipeline is designed to operate on input annotatables in hg38-based coordinates and that this will be the genome that annotators use by default. Annotator 1 (an “effect_annotator”) outputs two attributes—“worst_effect,” the worst predicted consequence across all affected gene transcripts, and “gene_list,” the list of genes affected by the input annotatable. Annotator 2 (a “position_score_annotator”) demonstrates a shorthand one-line annotator definition. This annotator generates a phyloP7way attribute that shows the site-wise evolutionary conservation rate across 7 vertebrate genomes at positions affected by the annotatable. The shorthand mode uses the implicit default annotation configuration associated with the resource, which, in this case, is to copy the only available score, phyloP7way, as an attribute. Annotator 3 (a “normalize_allele_annotator”) produces a “normalized_allele” (the implicit default name for this attribute) for downstream allele-level queries. Annotator 4 (an “allele_score_annotator”) queries the gnomAD (v4.1.0) resource using the “normalized_allele” and records only the AF (alternate-allele frequency) score, renaming it to “gnomAD_4.1_af”, capturing the resource version. Annotator 5 (an “allele_score_annotator”) also takes the “normalized_allele” and extracts the “MVP_rankscore^56^” from the “dbNSFP v4.9a” resource. Although dbNSFP aggregates more than four hundred scores, this pipeline selects only the “MVP_rankscore”. If needed, a user can trivially extract any of the other scores dbNSFP^38^ provides. Annotator 6 (an “allele_score_annotator”) also uses the “normalized_allele” and returns two scores from the ClinVar^19^ resource downloaded on 10/19/25 (“hg38/scores/ClinVar_20251019”): the “CLNSIG” score representing the variant’s clinical significance (e.g., benign/pathogenic) and “CLNDN” providing the associated disease term. Annotator 7 (a “gene_score_annotator”) uses the list of genes affected by the annotatable (available as the “gene_list” attribute produced by Annotator 1) and outputs the pLI_rank values for these genes from the “gene_properties/gene_scores/pLI” resource. Annotator 8 (a “gene_set_annotator”) uses the “gene_properties/gene_sets/MSigDB_curated/7.5” resource together with the available “gene_list” attribute to add a true-or-false attribute “KEGG_APOPTOSIS” indicating whether the genes affected by the annotatable belong to the corresponding MSigDB pathway^52^. Annotator 9 (a “liftover_annotator”) applies the “hg38_to_hg19” liftover chain resource to the default “input_annotatable” to produce the internal “hg19_annotatable” attribute that represents the “input_annotatable” in the coordinate space of the hg19 reference genome, and Annotator 10 (an “allele_score_annotator”) uses the “hg19_annotatable” to retrieve MPC (Missense badness, PolyPhen-2, and Constraint) from the hg19-based “hg19/scores/MPC” resource. Finally, the definition of annotator 11 demonstrates a powerful feature of GAIn’s pipelines: multiple annotators can be described in a single line in the annotation pipeline using resource pattern matching. In this example, the line represents multiple position_score_annotators, one for each resource that matches the pattern “TF*[target=CREB*]”, meaning that (i) the resource ID starts with “TF” corresponding to the transcription factor ChIP-seq resources from our GRR-ENCODE and (ii) the resource has a label called “target” with values starting with CREB. This pattern matches 19 ENCODE transcription-factor ChIP-seq resources targeting CREB1, CREB3, CREB5, CREBL2, and CREBBP.

GAIn pipelines can be added to a GRR as a resource of type “annotation_pipeline”. The summary page for an “annotation_pipeline” resource contains a description of the pipeline, including the generated attributes with detailed descriptions and, where appropriate, value histograms, as well as links to the summary pages of the underlying resources and to documentation for the annotators used by the pipeline. For example, the clinical annotation with ID “pipeline/hg38_clinical_annotation” in our main GRR includes 13 annotators covering variant effect, clinical significance, mutation damage, pathogenicity, allele frequency, and mutation tolerance, and is documented on its GRR summary page (https://grr.iossifovlab.com/pipeline/hg38_clinical_annotation).

#### GAIn Web Interface

GAIn’s web interface provides easy access to the system’s functionality without local setup (Figure 6). The web interface is connected to our main GRR and GRR-ENCODE. For annotation, users may either select from available annotation pipelines, which cover a wide range of annotation options, or create their own pipelines using the integrated pipeline authoring tool. Detailed descriptions of these features are provided in the “Getting started on the web” and “GAIn web interface” sections of the GAIn documentation.

**Figure 6.**
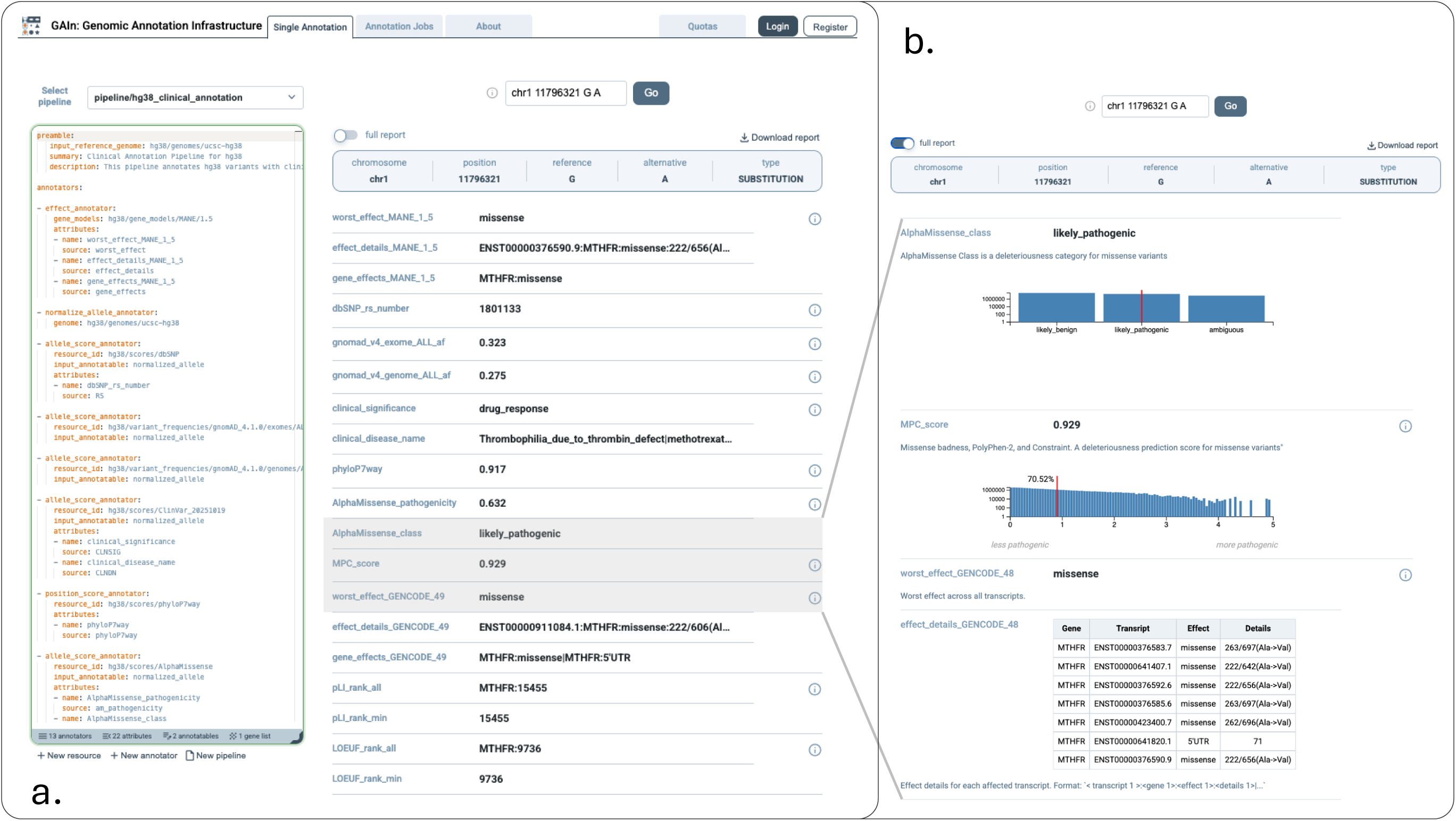
Example usage of Single Annotation in the GAIn web interface. **(a)** Left, the annotation pipeline editor, where users can select a previously saved pipeline or build a custom pipeline by defining the preamble and annotators. Right, example output for a single-allele query with compact report enabled, showing only the annotation attributes. The displayed attributes include effect details, gnomAD v4.1 allele frequency, AlphaMissense pathogenicity, clinical significance, dbSNP identifier, and LOEUF rank. **(b)** Example output for the same query with compact report disabled. In this view, the annotation results are shown together with additional explanatory text and, where available, score distributions. Histograms are shown in blue, and the score of the queried variant is indicated by a red vertical line.

GAIn supports two annotation modes. In “Single Annotation” mode, users provide a single annotatable (variant, position, or region) and either select a published pipeline available in our GRR or create a custom pipeline. The annotation runs immediately, showing the values of the annotation attributes that the pipeline produces for the selected annotatable, along with detailed descriptions for each attribute, including the resources used, value ranges, and histograms, where appropriate. This mode is ideal for rapid prototyping and pipeline validation before using these pipelines for large annotation jobs. In “Annotation Jobs” mode, users upload an input file, such as a TSV or VCF, choose an existing pipeline or provide a new one, and submit the job for server-side execution on GAIn’s computational infrastructure. The interface allows users to monitor job progress and download the completed results when processing finishes. For reproducibility, each job is associated with the exact setup used for the run, including the annotation pipeline. Optional user accounts provide access to the history of single-annotation alleles and annotation jobs, as well as storage for custom annotation pipelines.

#### GAIn Command Line Tools

GAIn is especially powerful on the command line: users can point to a public or private GRR and annotate millions of variants, positions, and regions while keeping pipelines and resources pinned for reproducibility. Below, we summarize the most frequently used tools. The “Getting Started on CLI” and “Annotation Infrastructure” sections of the GAIn documentation provide detailed usage instructions.

**“grr_browse”** lists the configured GRRs and allows listing and searching of the accessible resources.

**“grr_manage”** manages custom GRRs (see below for a technical description of how custom GRRs are built). The “grr_manage” tool verifies that all resources are properly configured, runs exhaustive validation tests for each resource, and produces an HTML summary that includes type-appropriate global statistics for each resource. For example, the summary lists the chromosomes and their lengths for reference genomes, the number of genes and gene transcripts for gene models resources, and score histograms for gene-, position-, and allele-score resources. Finally, the tool enables version control of custom GRRs using Git and DVC^57^, with support for large files commonly used as genomic resources.

**“annotate_doc”** renders human-readable documentation for an annotation pipeline, summarizing annotator types, referenced resources, and the set of output attributes (with basic distribution summaries where available). Generating pipeline documentation is a powerful way to validate it.

The tools above prepare for annotation—confirming resources and validating the pipeline. The tools below perform annotation given input files containing annotatables (Figure 1 “user input”) and a pipeline.

**“annotate_tabular”** applies an annotation pipeline defined in a YAML file to a tabular file with one annotatable per line. The annotatable type is controlled through command-line parameters and the available columns in the file. The tool produces a new table with the same number of lines as the input table, including all the columns from the input table, and appends one additional column for each attribute generated by the annotation process.

**“annotate_vcf”** applies an annotation pipeline to an input VCF (variant call format) file and produces an output VCF file in which annotation attributes are emitted as new INFO fields with corresponding header lines.

“grr_manage”, “annotate_tabular”, and “annotate_vcf” support parallel execution on multicore hosts and on computation clusters such as Sun Grid Engine (SGE) or SLURM. These parallelization features allow users to seamlessly leverage available computational infrastructure to manage large GRRs and to annotate hundreds of millions of annotatables.

#### Reannotation

Annotation pipelines change because new resources or resource versions become available (e.g., new gnomAD releases, updated gene models, refreshed conservation tracks). Often, only a small part of the pipeline, such as a single annotator, needs to be changed. In such cases, updating the annotations of large datasets (e.g., tens of millions of records typical in large genomic projects) with the entire pipeline can be wasteful. GAIn’s re-annotation feature identifies which annotators changed between the old and new annotation pipelines and runs only the modified annotators, saving considerable computation time and costs in genomic annotations across large projects. In practice, users point GAIn to the results of a prior annotation, the pipeline that was used, and the desired new annotation pipeline, and GAIn produces revised annotation results. This approach enables routine, low-friction updates as resources and requirements evolve.

### GAIn extensions

#### GRR Configuration

When freshly installed, GAIn uses resources from our main GRR by default. Users who need additional resources can create their own GRR (see below) and configure GAIn to search multiple GRRs in a specified order, including our main GRR, GRR-ENCODE, other public GRRs, and any number of personal GRRs. This configuration is specified in a YAML file in the user’s home directory named “.grr_definition.yaml,” whose structure and usage are described in detail in GAIn’s documentation.

GAIn also supports optional caching of remote GRRs. When caching is enabled, and a resource is requested from a remote GRR, GAIn first downloads the resource into a local cache directory and then uses the cached copy for annotation. Caching is essential for throughput when annotating large datasets, but it requires sufficient local disk space and time for the initial downloads, because some resources can be tens of gigabytes in size (file sizes are listed on each resource’s information page).

#### Custom GRRs

Creating a local private GRR is relatively straightforward. A local GRR is simply a directory on disk that GAIn treats as a catalog of resources. Each subdirectory at any level inside the GRR directory that contains a genomic_resource.yaml file represents a genomic resource. The path from the GRR directory to the resource directory is treated as the resource ID. The genomic_resource.yaml file specifies the resource configuration, including the resource type, the associated data files and their formats, and a description of the resource, such as its source and version. The associated files are stored in the resource directory, which may also contain scripts for downloading and transforming resource files from the original source.

Users can create an empty repository with “grr_manage init” and populate it with the required resources by creating the sub-directory structure and the genomic_resource.yaml files. They can use the additional functions (resource-repair and repo-repair) of the “grr_manage” tool to validate the GRR and create resource statistics and information pages for the configured resources. GAIn also supports version control of GRRs using Git together with DVC^57^ for large resource files. Users may also choose to make their local GRRs publicly available to facilitate reuse by others.

We have prepared two resources to help new users create their own GRRs. First, GAIn’s documentation includes a substantial “Getting Started with GRRs” guide in addition to the detailed description of the relevant concepts and tools. Second, we provide mini-GRR, a small, self-contained Genomic Resource Repository on GitHub (https://github.com/iossifovlab/mini-GRR) that includes example configurations for many resource types and file formats. mini-GRR includes a toy genome (two short chromosomes, 20 nucleotides each), examples of gene models (RefSeq- and GTF-style, one gene per chromosome), position scores, allele scores, gene scores, gene sets, and CNV collections. Resources span common file types (e.g., TSV/tabix, BedGraph/BigWig, VCF) and coordinate conventions (0-based and 1-based), making explicit how formats and offsets are handled.

In their private GRRs, users can include all resource types supported by GAIn (see above), including reference genomes, gene models, and position scores. These resources can be publicly available resources that are not yet included in any public GRR or, importantly, private resources generated by users.

#### Custom Annotators

GAIn supports extension through a Python-based plugin architecture. GAIn plugins are implemented as Python packages and can add new functionality, including new annotator types. A custom annotator can operate on its input annotatable, query genomic resources through GAIn’s resource access API, and use attributes generated by annotators earlier in the pipeline to compute new attributes. For example, a custom annotator can combine effect predictions, conservation scores, and allele frequencies into a single prioritization flag indicating whether the annotatable merits further experimental follow-up.

Here, we illustrate this with two concrete plugin examples that we have implemented. First, we implemented a “vep_gtf_annotator” plugin that delegates consequence prediction to the widely used Ensembl Variant Effect Predictor (VEP)^22^, allowing GAIn pipelines to incorporate VEP-derived annotations alongside other native GAIn annotations. Second, we implemented a “spliceai_annotator” plugin that predicts the splice-altering effects of variants using the SpliceAI deep learning model^58^. Instead of relying on precomputed genome-wide score tables, it computes splice-impact scores on the fly from the variant and reference sequences using a pre-trained deep neural network. The “spliceai_annotator” depends on TensorFlow^59^ and, although it works correctly on CPUs, is computationally intensive and is therefore primarily intended to run on hosts with available GPUs.

## Discussion

GAIn addresses a practical bottleneck in genomic interpretation. Although many powerful annotation resources already exist, they are distributed across different file formats, genome assemblies, versions, and coordinate conventions, making them difficult to combine and maintain consistently. In this work, we present GAIn as a framework for genomic annotation. Its central contribution is to make annotation workflows more transparent, reproducible, and maintainable by combining declarative pipelines with curated Genomic Resource Repositories (GRRs). By separating workflow logic from resource management, GAIn makes it easier to organize resources, preserve provenance, reuse annotations across assemblies, and build composable workflows that can be updated over time. This explicit separation is also important scientifically, because version-pinned resources and declarative pipelines make annotation results easier to audit, compare, and revisit.

A major strength of GAIn is that it supports multiple annotation use cases within a single framework. In contrast to many existing systems that primarily focus on variant interpretation, GAIn can annotate genomic variants, positions, and regions. This broader scope is important because modern studies increasingly need to interpret not only sequence variation, but also genomic signals such as GWAS hits, QTLs, ChIP-seq peaks, ATAC-seq peaks, and other intervals.

GAIn is designed to support both exploratory and large-scale use. The web interface enables rapid testing and interactive inspection of annotations, whereas the command-line tools support large studies with hundreds of millions of variants through easy-to-use and flexible parallelization. This combination is especially useful for groups that need to rerun analyses over time as resources, assemblies, and study-specific requirements change. In addition, selective re-annotation provides an important practical advantage by allowing users to update only the affected parts of an annotation workflow when resources change, rather than rerunning entire pipelines. GAIn also makes it straightforward to use resources defined in reference assemblies that do not match the input assembly through liftover, enabling rich annotations even for projects using new assemblies such as T2T-CHM13/hs1. More broadly, this helps prevent analyses from being locked to a single reference assembly and extends the usable life of existing resources.

Our public Genomic Resource Repositories (main GRR and GRR-ENCODE) lower the barrier to entry by providing immediately usable, curated resources, while custom GRRs allow individual groups to incorporate local, private, or domain-specific datasets. The separate GRR-ENCODE repository demonstrates that the same framework can also accommodate very large, specialized collections without compromising the usability of the main repository. We hope that investigators who develop new genomic resources will find GAIn’s GRR management features useful for packaging their resources as GRRs and, when appropriate, making them publicly available for others to use. At the same time, the plugin architecture makes GAIn extensible, enabling new annotators and external methods to be integrated without modifying the core system.

Machine-learning-based function predictors are becoming increasingly common in variant interpretation. GAIn’s plugin architecture provides a direct way to integrate such models into annotation pipelines, either by running a model at query time (as in our spliceai_annotator) or by managing precomputed predictions as versioned resources. In addition, the GRR abstraction can help organize and track the large datasets used to train and evaluate these models, together with their provenance and versions.

Future work can expand the public repositories, support additional annotator types and resource classes, and improve interoperability with external methods and downstream analysis environments. In its current state, GAIn provides a practical, end-to-end framework for variant annotation by combining curated GRRs, declarative pipelines, and multiple execution modes (command-line and web), while remaining extensible via plugins. As a result, GAIn provides a foundation for maintaining reusable annotation workflows over time, while leaving a clear path for continued expansion of resources and capabilities.

## Notes

### Competing Interest Statement

The authors declare no competing interests.

